# An Evaluation of Physiological Responses Towards Variant Respiratory Function

**DOI:** 10.1101/2023.10.23.563450

**Authors:** Tianchen Wang, Alfred E. Mann

## Abstract

The respiratory system is listed among the most critical systems that sustain lives. Numerous studies have explored respiratory system mechanisms and developed models to understand its responses to external stimuli in recent years. However, the relationship between the function and other physiological appearances must be well-established. In this study, the respiratory function was evaluated as inspiratory capacity (IC), and various physiological responses were linked to the variance of such function. Paired t-tests confirmed significant inter-posture variations on IC. A positive linear correlation was observed between MEP and IC, while a weaker was observed between abdominal muscle shortening and IC, particularly during the end exhalation. The lung inflation reflex investigation noted a strong linear correlation between heart rate and breath level. This study could shed light on understanding the respiratory system and its applications in various fields.

## Introduction

The respiratory system is listed among the most critical systems that sustain lives. The system generally contains several key components, including the lungs, tracheas, heart, and various muscular and skeletal elements in the human body. It is a complex system with multiple feedback loops for facilitating gas exchanges in the alveoli and oxygen delivery to the rest of the body. Understanding the mechanism behind such a system would provide valuable aid in diagnosing and treating respiratory diseases and guiding people to maintain a healthy respiratory function. Numerous studies have explored respiratory system mechanisms and developed models to understand its responses to external stimuli in recent years. It’s known that respiratory response, breathing pattern every respiratory cycle, to exercise varies according to the intensity of exercise the subject receives (Serna, et al.).

Since the respiratory system takes input from multiple sources, respiratory function can be measured in several ways. Until now, such relationships between the function and other physiological appearances are not yet well-established. This experiment evaluated the function as inspiratory capacity (IC) and quantified by an incentive spirometer. The effect of different body postures on IC was verified with t-tests. Then, IC was linked to muscular activity by measuring the maximum expiratory pressure (MEP). The inflation reflex was also tested to evaluate the impact of breathing on the cardio system. The variance in body posture would significantly affect respiratory capability. A positive relationship between MEP and IC should be quantitively constructed regarding abdominal muscular activity. Due to an inflation reflex feedback loop, the cardio activity represented as the heart rate should increase as the subject increases the breathing rate compared to the resting level. The experiments were expected to draw connections between respiratory functions and various identities within/outside the respiratory system.

## Materials and Methods

### 1) Inspiratory Capacity (IC) Measurements

A Voldyne 5000 incentive spirometer was used to obtain the measurements of inspiratory capacity (IC). Before the measurement, a ruler with millimeter markings was attached to the spirometer for a more precise reading, where the zero mark on the ruler was aligned with the upper edge of the piston. The subject was told to sit upright on the chair, ensuring a clear visibility of the piston level. During the data collection, the subject inhaled through the tube connected to the spirometer while maintaining a flow rate within the “best” range marked on the cylinder. The maximum piston level was recorded in millimeters progressed along the ruler when the subject completed a full inhalation and exhaled normally, disconnected from the spirometer. After about a minute of recovery between each measurement, the process was repeated five times for two sets before the subject changed to a supine posture and underwent the same process. A total of 10 data points for each posture were recorded and converted to IC in body temperature, ambient pressure, and gas saturated with water vapor (BTPS) and standard temperature and pressure and dry (STPD) using the formula.

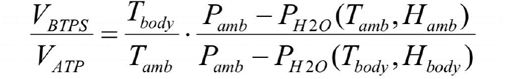

The averaged IC-BTPS measurements for each posture were compared with the predicted standard value using the nomogram.

### 2) Statistical Comparison

Building on those IC measurements, a statistical comparison involving t-tests was applied to evaluate the potential intra- and inter-group variability of the two different postures. The average differences and the standard deviation of such were calculated from the 20 paired comparisons between the seated and supine group in the treatment-to-treatment t-test, resulting in 10 paired differences. Test-to-test t-tests were set up with the same calculation after pooling all the measurements from the same group and formatting five pairs within each group. From that, the t statistic could be calculated and compared to the reference value of a specific degree of freedom and level of significance to see if the data points collected were minimally affected by the random variation nature of physiological measurements.

### 3) Maximum Expiratory Pressure (MEP)

MEP was measured using a blood pressure gauge with the rubber tube disconnected from the arm cuff. The subject was told to exhale into a gage inlet to measure MEP after taking a full breath (maximum IC) while seated with the nose closed and no airflow present. Data was recorded after the pressure was maintained for at least one second. The three highest pressures out of the four points were kept and averaged. Following a 5-minute rest between each set, the procedure was repeated with the subject inhaled to 2/3 and 1/3 of the maximum IC determined in **1)**. In addition, the abdominal length was roughly estimated by measuring the length of the subject’s abdomen for each measurement. The three maximum pressures and estimated abdominal muscle length measured in centimeters vs. IC volume were plotted for analysis in Excel.

### 4) Inflation Reflex

Inflation reflex on heart rate was assessed with an incentive spirometer to control inspired breath volumes ranging from normal (0.5 liters) to three times normal (1.5 liters). The resulting heart rate was measured with a smartphone app. The spirometer was placed on a table to obtain accurate measurements. After a 5-minute break between each set, the subject was told to increase the inhaling volume according to the scheme and refer to the readings on the spirometer. The heart rate vs. inspired breath volume was plotted for analysis in Excel.

## Data & Results

The raw data was collected from the lab and plugged into Excel for further analysis. The inspiratory capacity within each set of measurements was consistent, where most of the data fell within the range of 3000 mL to 3500 mL, and the averaged results for the two postures were around one-tenth greater than the reference value (Figure 1). The consistency between sets with the same posture and the variance across different postures was verified in the subsequent t-test analysis, where there was a far more significant difference from treatment to treatment when compared with the difference from test to test (Figure 2, Figure 3.1, Figure 3.2). A positive linear correlation between MEP and IC was identified within the measurement range (Figure 4). The linear relationship was present but could have been stronger between abdominal muscle length and IC, especially when the exhalation was nearly complete (Figure 5). In detecting the lung inflation reflex, a strong linear correlation was observed between heart rate and breath level (Figure 6).

**Figure 1.**
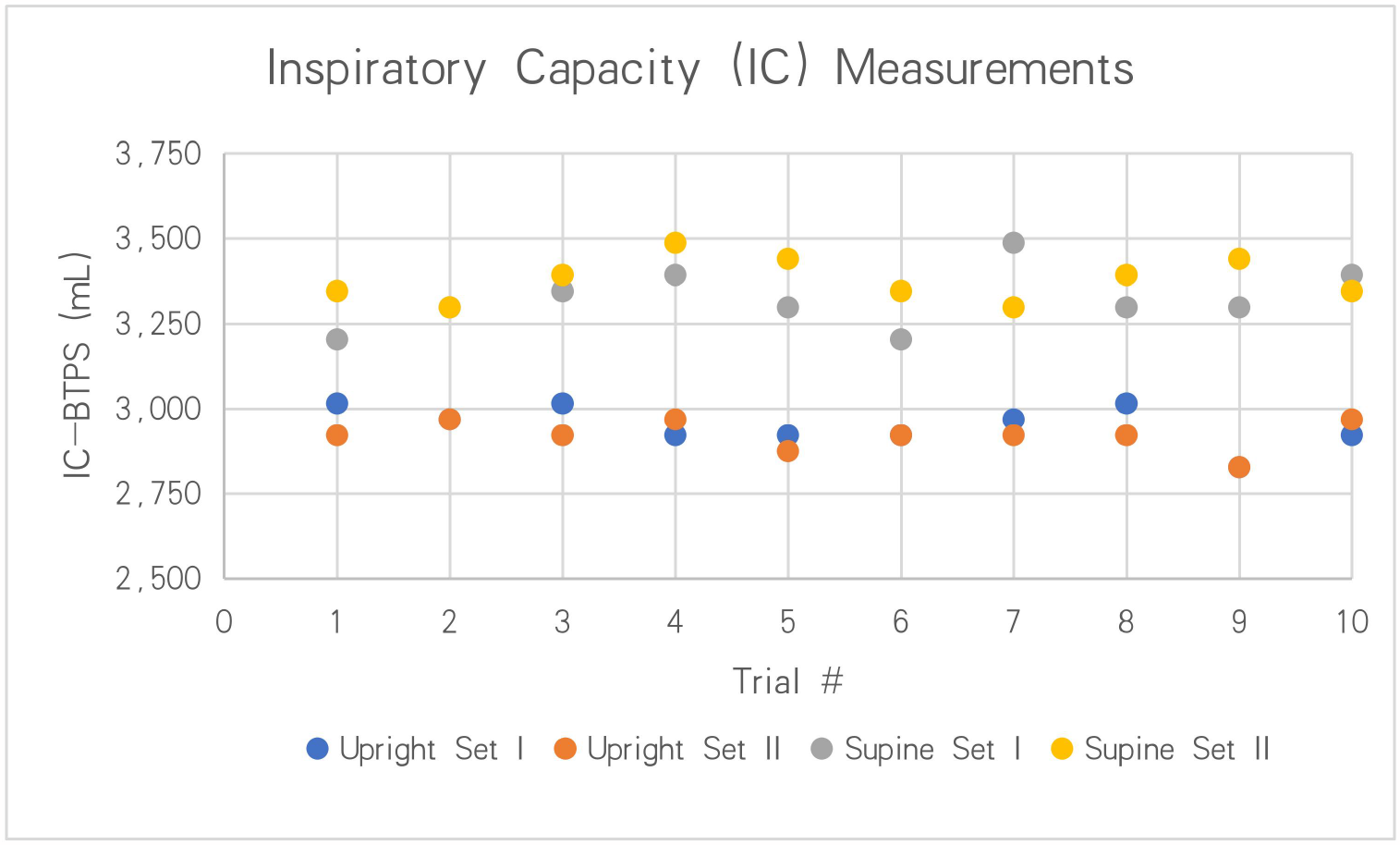
Inspiratory capacity (IC) measurements scatter plot. For each set of measurements, 10 data points were recorded. The millimeter readings were converted to IC-BTPS in milliliters. There was a one-minute break between each measurement and a five-minute break between each set. The upright and the supine posture was 102.06% and 116.90% greater than the reference value.

**Figure 2.**
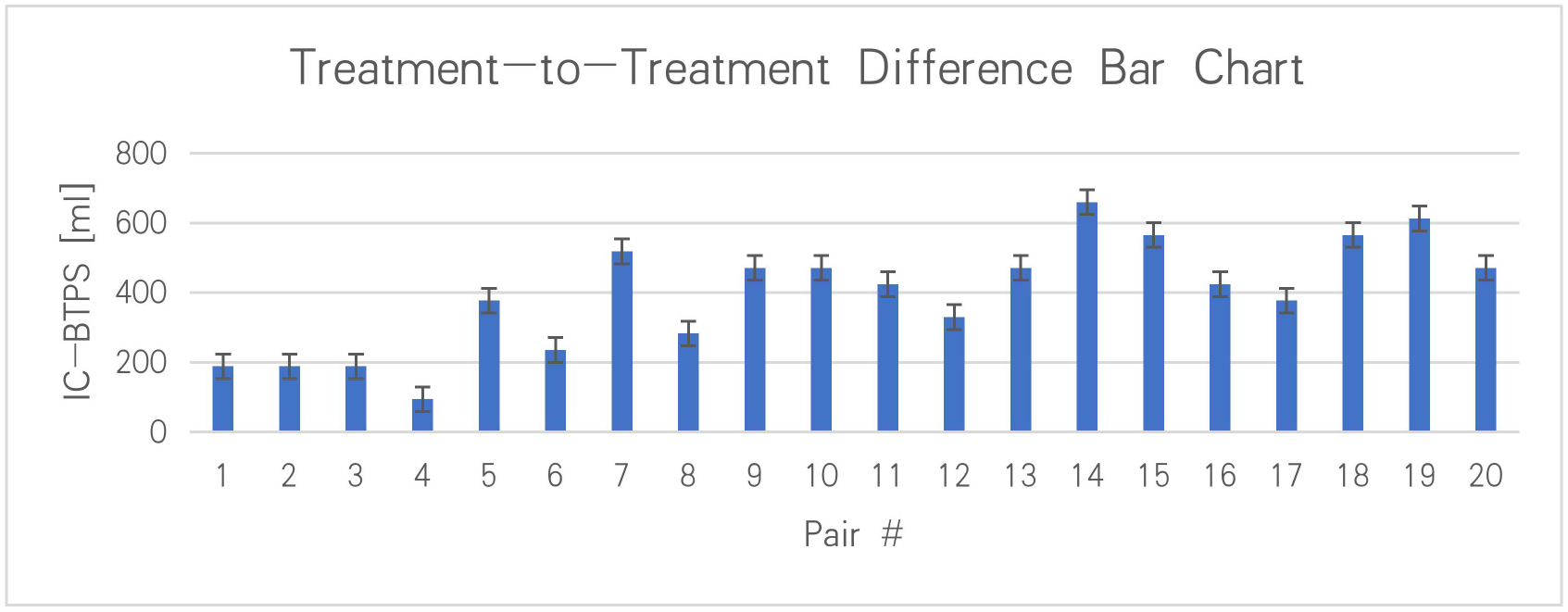
Treatment-to-Treatment Difference Bar Chart. Each difference was calculated by subtracting supine posture data from upright posture data. The 20 pairs were randomly formatted from the pooled data points collected in previous experiments. Error bars were generated +/- one standard deviation from the origin using sample size=20 and p-value=0.05. The resulting t-statistics=11.17 with a degree of freedom=19, p<0.0005.

**Figure 3.1.**
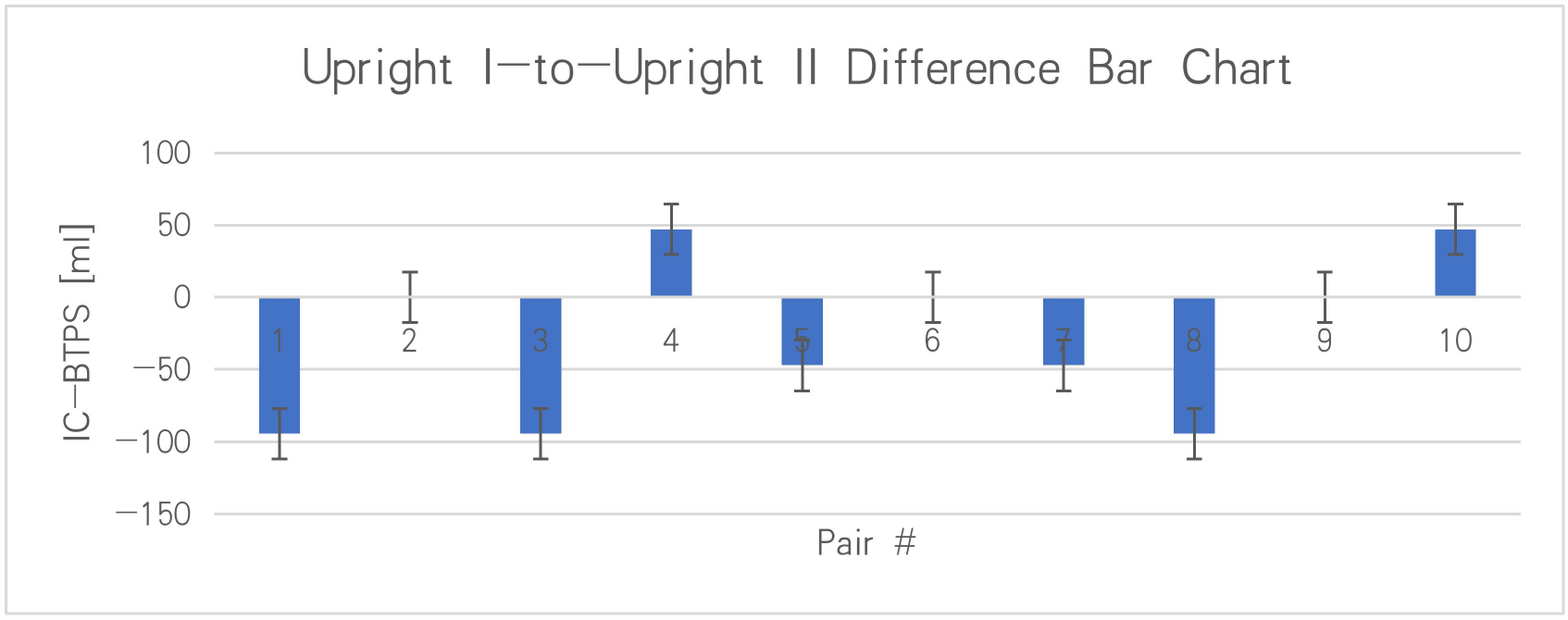
Upright Set I to Set II Difference Bar Chart. Each difference was calculated by subtracting supine posture data from upright posture data. The ten pairs were randomly formatted from the pooled data points collected in previous experiments. Error bars were generated +/- one standard deviation from the origin using sample size=20 and p-value=0.05. The resulting t-statistics=-1.62 with a degree of freedom=9, p>0.05.

**Figure 3.2.**
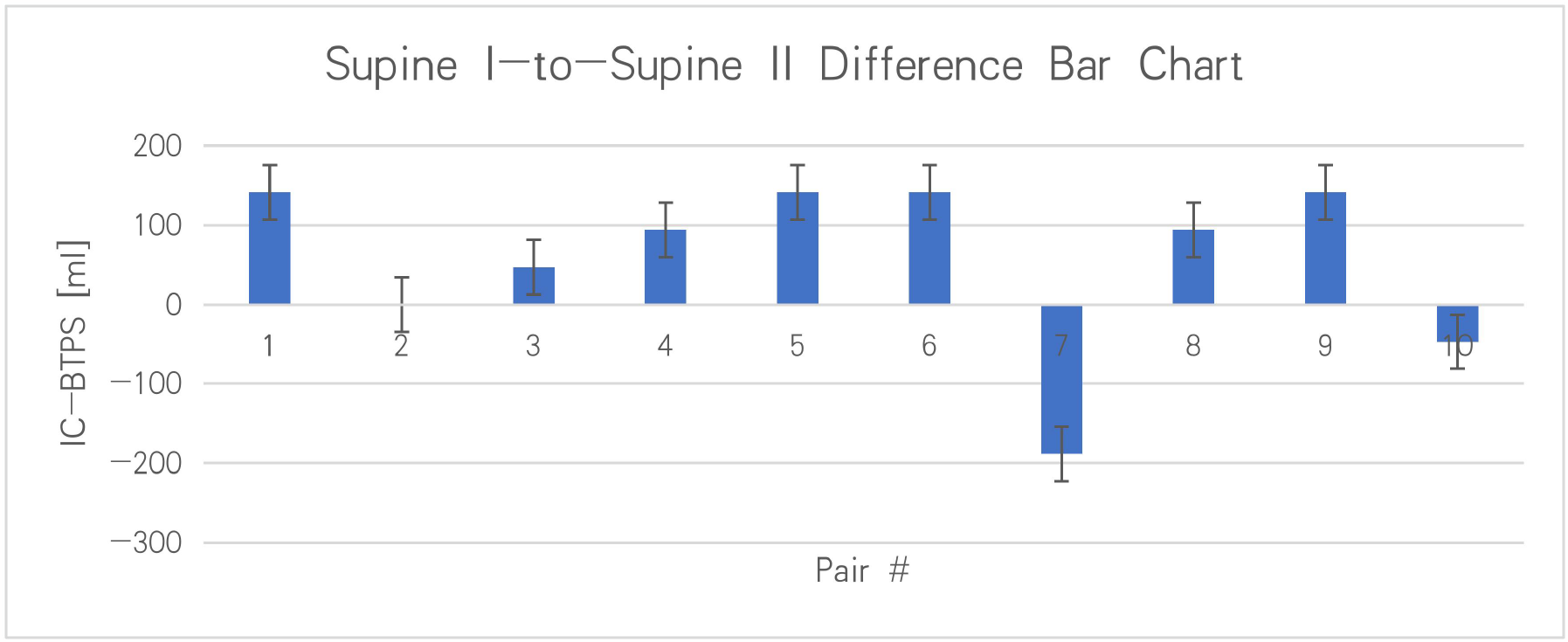
Supine Set I to Set II Difference Bar Chart. Each difference was calculated by subtracting supine posture data from upright posture data. The ten pairs were randomly formatted from the pooled data points collected in previous experiments. Error bars were generated +/- one standard deviation from the origin using sample size=20 and p-value=0.05. The resulting t-statistics=1.65 with a degree of freedom=9, p>0.05.

**Figure 4.**
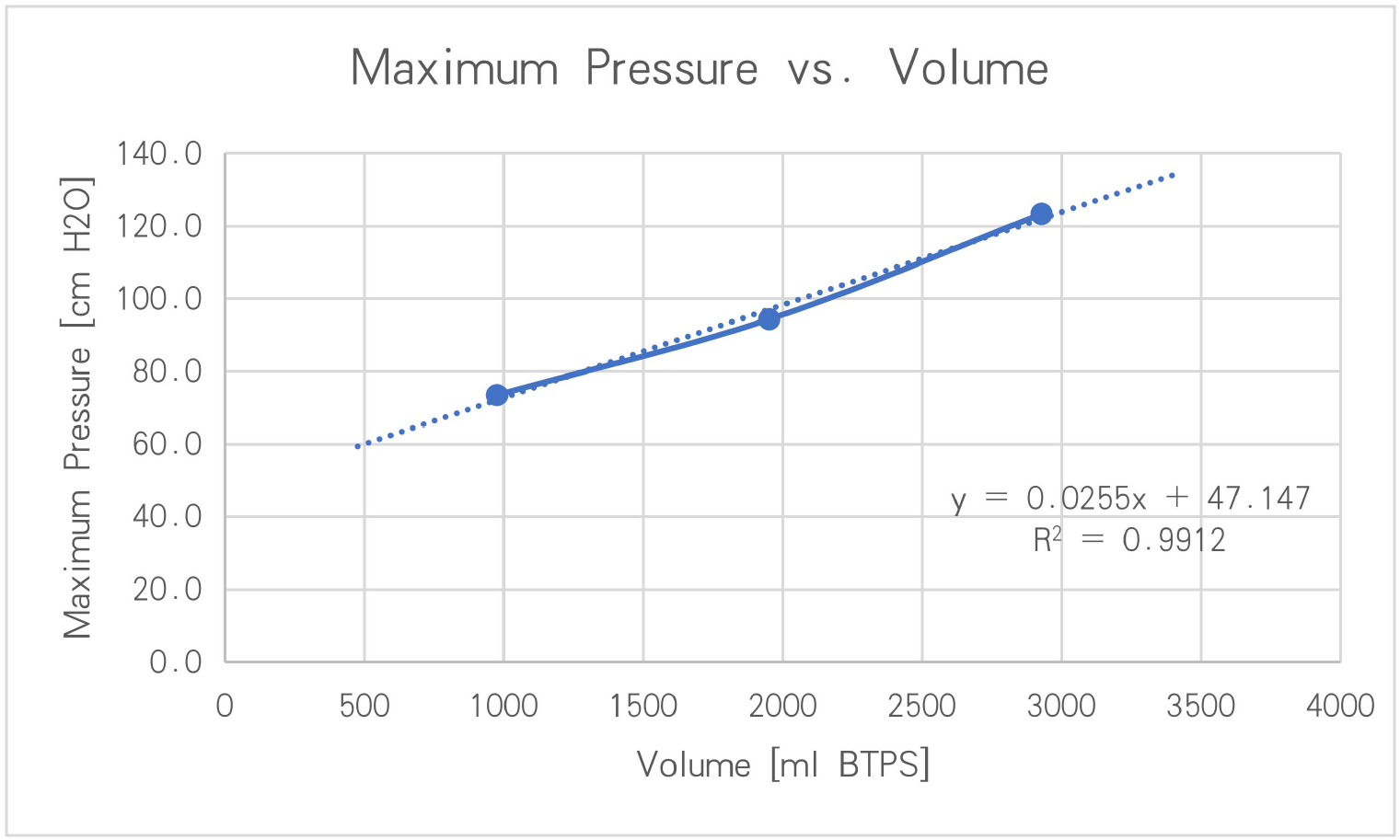
MEP vs. IC line plot. Data points were taken at three levels corresponding to maximum IC, 2/3 IC, and 1/3 IC. A line of best fit was generated using a linear regression model with an R^2^ value of 0.9912.

**Figure 5.**
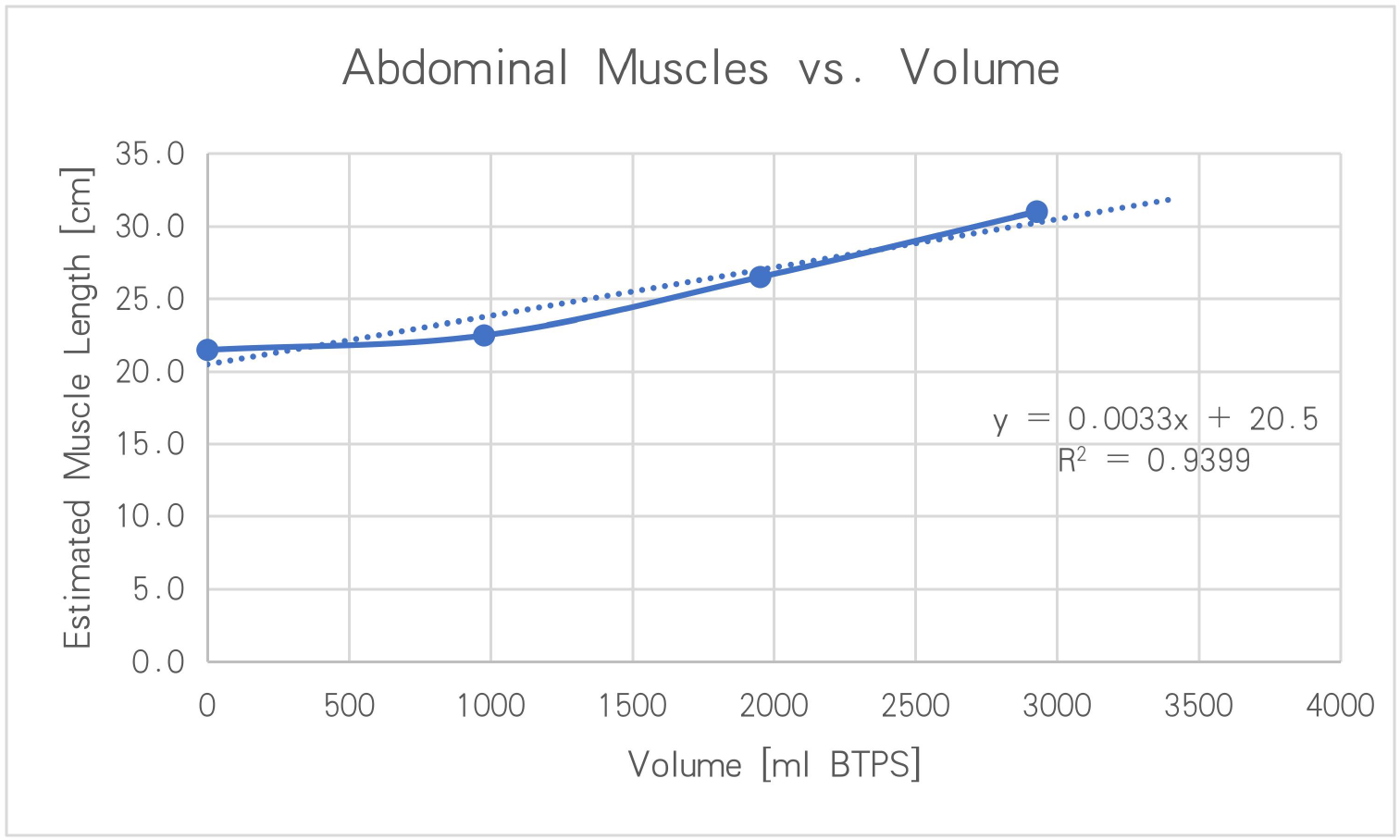
Estimated Abdominal Muscle vs. IC line plot. Data points were taken at four levels corresponding to maximum IC, 2/3 IC, 1/3 IC, and full expiration. A line of best fit was generated using a linear regression model with an R^2^ value of 0.9399.

**Figure 6.**
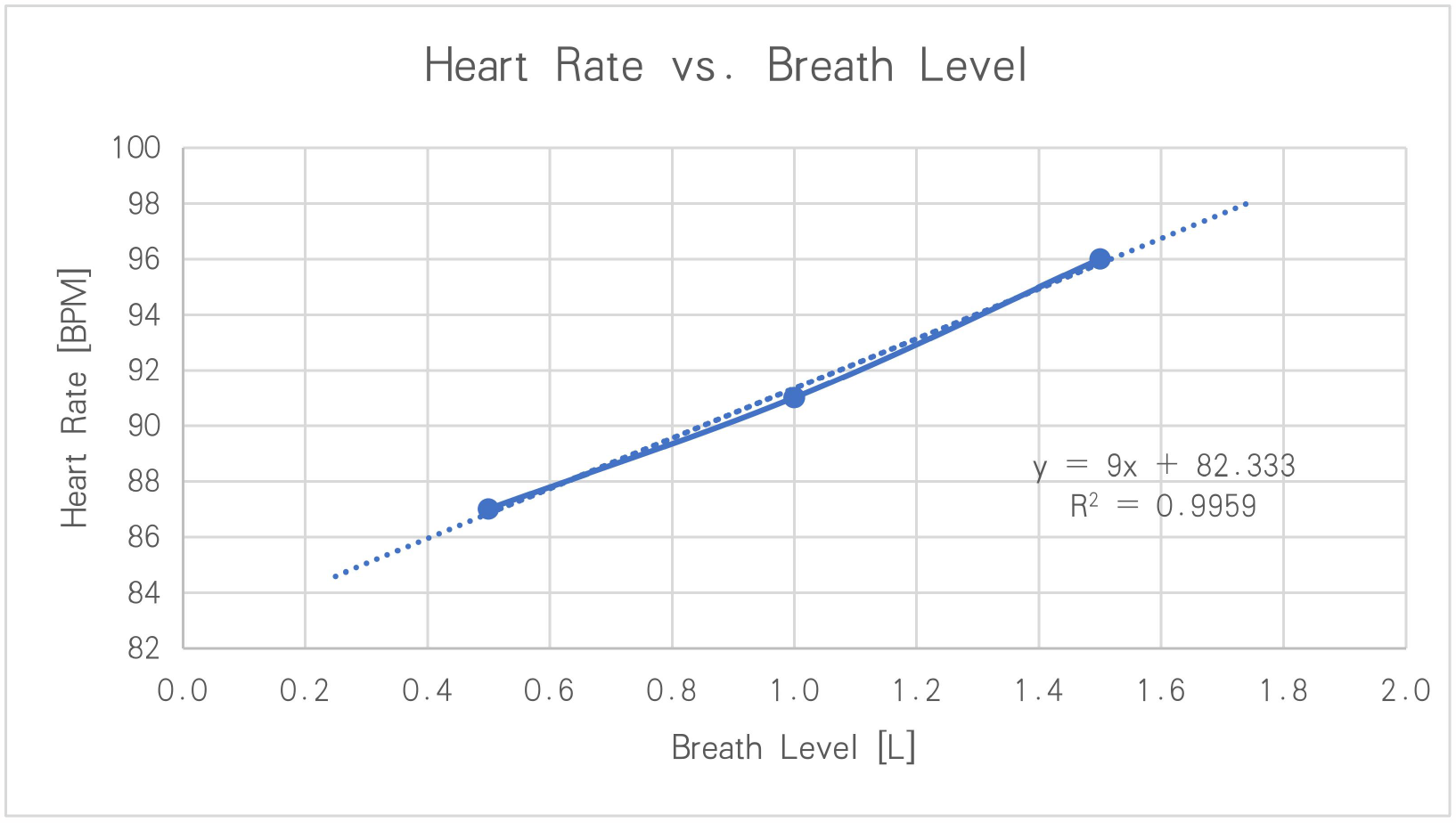
Heart Rate vs. Breath Level line plot. Data points were taken at three levels corresponding to 0.5L, 1.0L, and 1.5L inhalation volume. A line of best fit was generated using a linear regression model with an R^2^ value of 0.9959.

## Discussion and Conclusion

The rationale behind using the paired t-tests on the pool of inspiratory capacity (IC) measurements is that physiological measurements come with variances, and eliminating such would be crucial for a practical evaluation of these data points in subsequent analysis. It is when paired t-tests became very useful in comparing two sets of data where each data point in one group is directly related to another. Based on the data collected, the paired treatment-to-treatment t-test determined a significant difference between the means of IC gathered from upright and supine postures, with a possibility of less than 0.05% that the differences were caused by randomness. The significant difference across the group means that the posture change could affect one’s ability to inhale air as much as possible. In addition, the paired test-to-test t-test confirmed that there was no significant difference between sets of the same posture, which explained the slight differences between them were solely contributed by randomness. The measurements were consistent and could be pooled into one common group.

Observing the maximum expiration pressure (MEP) vs. IC, a robust linear relationship could be confirmed as the R^2^ value was close to one. Combined with part of the Abdominal Muscle Length vs. IC plot, the active role of the abdominal muscle in participating in the respiratory mechanism could be verified. MEP increase brought by the shortening of the abdominal muscle was evident, especially when the subject inhaled a large volume of air and expired rapidly. Similar roles of the abdominal muscle have been reported in other papers. The abdominal expiratory muscles were significantly thicker in patients with simple weaning than those with difficult or prolonged weaning (Amara et al.). The diminishing effect of abdominal shortening in increasing MEP as the subject inhaled a smaller air volume suggests that components other than the abdominal muscle could contribute to the expiration process. The reflex cardiovascular responses to lung inflation were identified as early as in the 1990s, where active lung inflation by a single deep inspiration in man gives rise to remarkable pressor response (Looga). Here, the linearly increased heart rate when the subject actively increased the inhalation volume verified the presence of such a feedback loop in respiratory mechanism regulation within the measured range from normal inhalation to maximal inhalation again.

Regarding limitations to this study, the study was constrained to only one subject, and some measurement ranges needed to be more comprehensive. For example, the data points taken in MEP vs. IC were 1/3 IC away from each other, which was a relatively huge gap. Similar confinements on the range of data appeared in the heart rate vs. breath level plot, where the effect of the inflation reflex was not evaluated when the subject was breathing below the resting level of breath volume. To conduct future experiments that expand on these findings, more subjects of different ages, sexes, and races could be recruited, and a more comprehensive range of measurements should be considered to understand those physiological responses to the varying respiratory function thoroughly. Built on the published study, which reported differences in the MEP during induced cough between the group who failed extubation and those without extubation failure (Carrera et al.), a comparison between those responses under pathological and normal conditions could also be added.

With careful regulation, the respiratory system’s function extends well beyond breathing. It plays crucial roles in other metabolic activities, such as maintaining the homeostasis of the blood circulation system and reacting to immune responses. As reported, SARS-CoV-2 predominantly affects the lungs but can also affect other organs (Chiner-Vives, 2022). Thus, understanding the interactions among physiological responses to respiratory function remains crucial. This study could shed light on various fields, from life-saving emergencies to maintaining a healthier life, and provoke more scientific research into the complex respiratory system.

